## Supplementary Figures S1-S11. Pre-registered behavioral results, replication experiment, model simulations, and MEG experiment supporting analyses for "Humans construct idiosyncratic, self-consistent global rankings from few-shot local evidence"

### S1 Figure

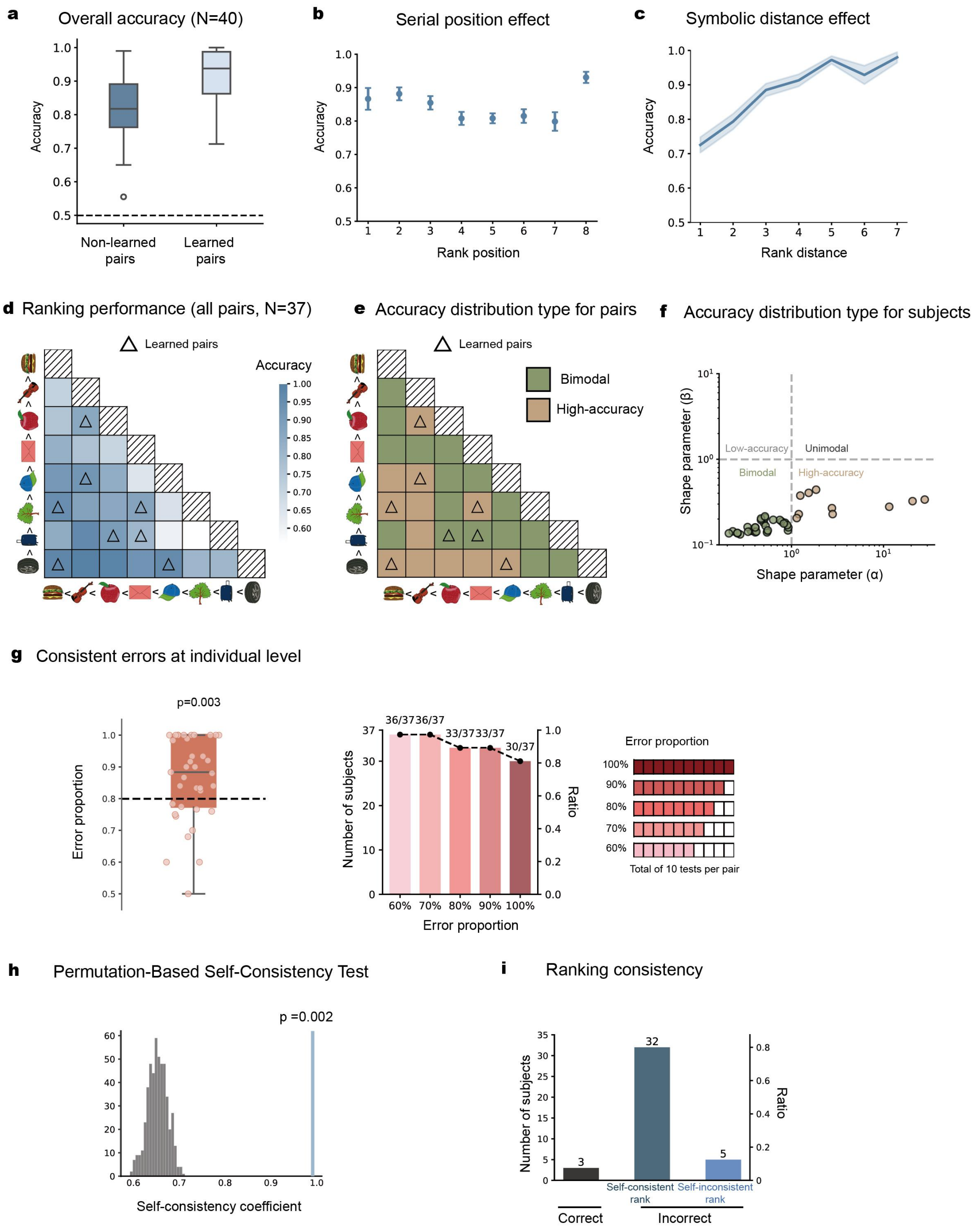


**S1 Fig. Pre-registered behavioral results (N = 40). (a)** Grand averaged ranking accuracy for learned (dark blue; 8 pairs in few-shot learning) and non-learned pairs (light blue; remaining 20 pairs not directly learned). **(b)** Grand averaged accuracy as function of rank position (serial position effect). **(c)** Grand averaged accuracy as function of ranking distance (distance effect). **(d)** Grand averaged accuracy matrix for all the 24 tested pairs, with row and column denoting corresponding items per pair (Deep-to-light color represent high-to-low accuracy). Eight directly-learned pairs (few-shot learning) were marked by triangles. **(e)** Beta-distribution model fitting results for each pair (brown: high-accuracy, $\alpha > 1$, $\beta < 1$; green; bimodal distribution, $\alpha < 1$, $\beta < 1$. **(f)** Beta-distribution model fitting results for each subject. Each point represents the best-fitting $\alpha$ and $\beta$ parameters for an individual subject. Brown points indicate subjects with bimodal error distributions ($\alpha < 1$, $\beta < 1$); green points indicate subjects exhibiting high-accuracy patterns ($\alpha > 1$, $\beta < 1$). Both $\alpha$ and $\beta$ are significantly smaller than 1 (one-sample *t* test, excluding 10 high accuracy participant as pre-registered; $\alpha$, $t(29) = -12.32$, $p < 0.001$; $\beta$, $t(29) = -185.69$, $p < 0.001$). **(g)** Individual-level error consistency (left) and proportion of subjects making consistent error on at least one local pair for different thresholds (0.6 to 1.0). Dots denote individual data. **(h)** Self-consistency was significantly higher than chance (permutation test: p = 0.002; observed = 0.99). **(i)** Number of subjects for each self-consistency category.

### S2 Figure

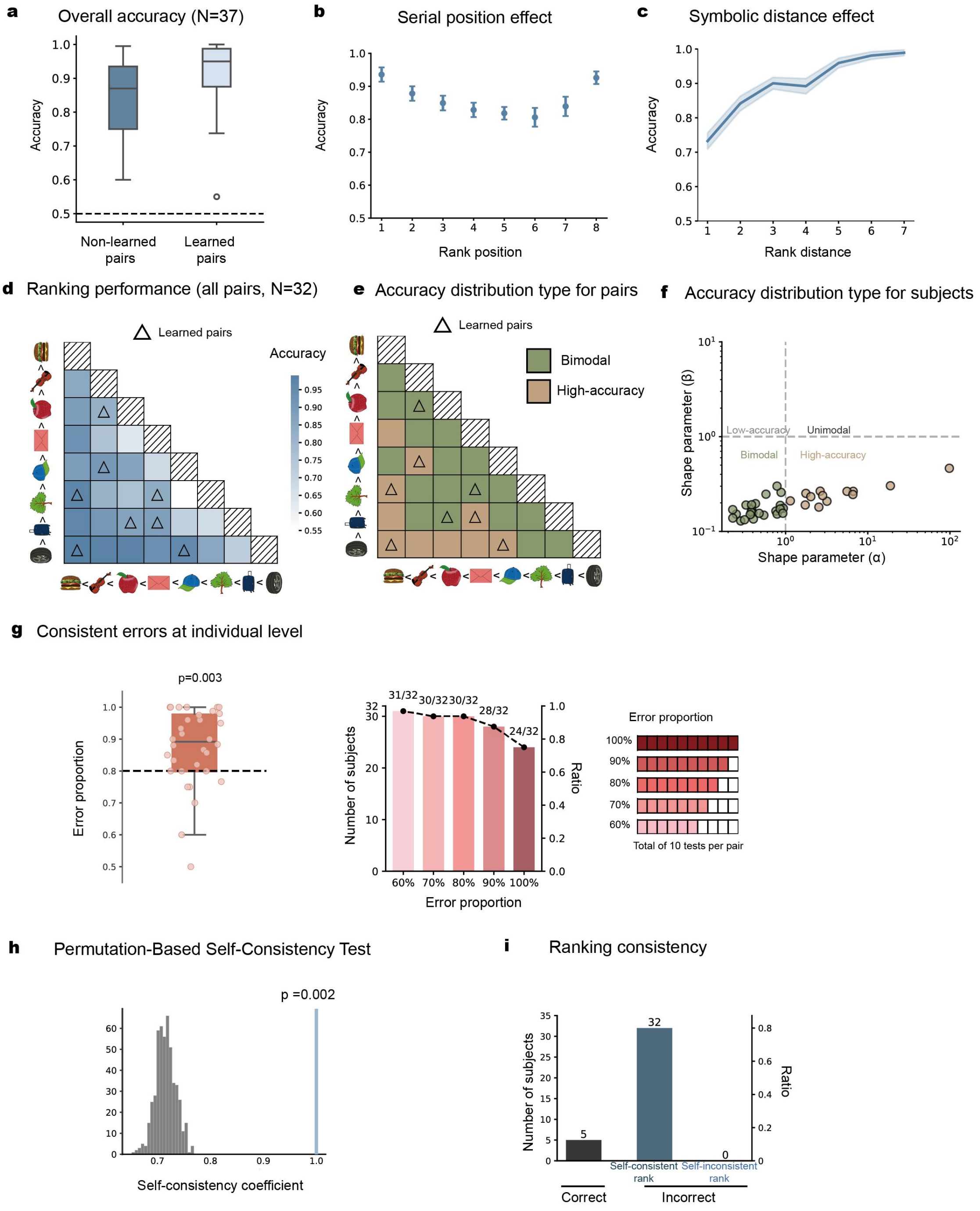


**S2 Fig. Replication behavioral results (*N* = 37). (a)** Grand averaged ranking accuracy for learned (dark blue; 8 pairs in few-shot learning) and non-learned pairs (light blue; remaining 20 pairs not directly learned). **(b)** Grand averaged accuracy as function of rank position (serial position effect). **(c)** Grand averaged accuracy as function of ranking distance (distance effect). **(d)** Grand averaged accuracy matrix for all the 24 tested pairs, with row and column denoting corresponding items per pair (Deep-to-light color represent high-to-low accuracy). Eight directly-learned pairs (few-shot learning) were marked by triangles. **(e)** Beta-distribution model fitting results for each pair (brown: high-accuracy, $\alpha > 1$, $\beta < 1$; green; bimodal distribution, $\alpha < 1$, $\beta < 1$. **(f)** Beta-distribution model fitting results for each subject. Each point represents the best-fitting α and β parameters for an individual subject. Brown points indicate subjects with bimodal error distributions ($\alpha < 1$, $\beta < 1$); green points indicate subjects exhibiting high-accuracy patterns ($\alpha > 1$, $\beta < 1$). Both α and β are significantly smaller than 1 (one-sample *t* test, excluding 13 high accuracy participant as pre-registered; α, $t(23) = -9.27$, $p < 0.001$; β, $t(23) = -97.30$, $p < 0.001$). **(g)** Individual-level error consistency (left) and proportion of subjects making consistent error on at least one local pair for different thresholds (0.6 to 1.0). Dots denote individual data. **(h)** Self-consistency was significantly higher than chance (permutation test: p = 0.002; observed = 1.00). **(i)** Number of subjects for each self-consistency category.

### S3 Figure

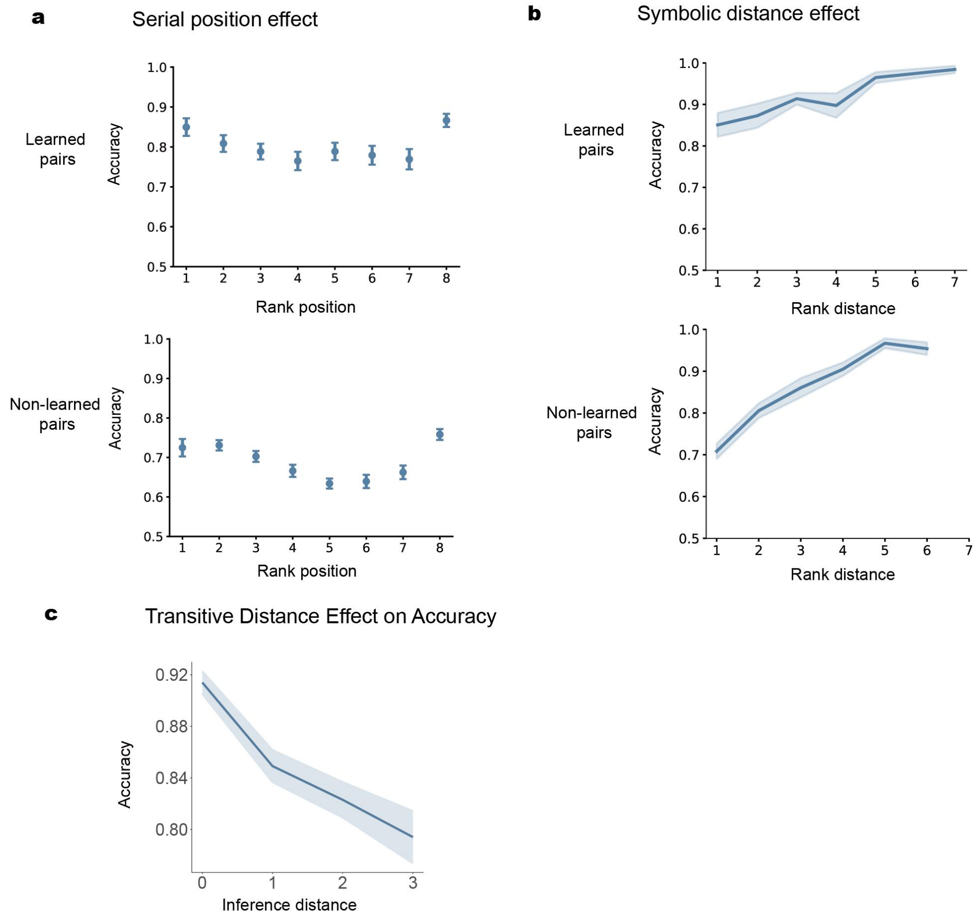


**S3 Fig. Serial position effect, distance effect for learned and non-learned pair. (a)** Serial position effect. Upper panel (learned): $F(7, 532) = 11.87$, $p < 0.001$; lower panel (non-learned): $F(7, 532) = 11.87$, $p < 0.001$. **(b)** Symbolic distance effect. Upper panel (learned): slope coefficient = 0.02, $t(76) = 4.556$, $p < 0.001$; lower panel (non-learned): slope coefficient = 0.05, $t(76) = 14.141$, $p < 0.001$. **(c)** Transitive distance effect is significant without controlling for rank distance linear mixed-effect model; $F(1, 1463.00)=6.37$, $p = 0.012$) but not significant after control ($F(1, 1463.00)= 0.17$, $p = 0.68$). Pairs with 0 transitive distance is excluded from this analysis as they are directly learned pairs.

### S4 Figure

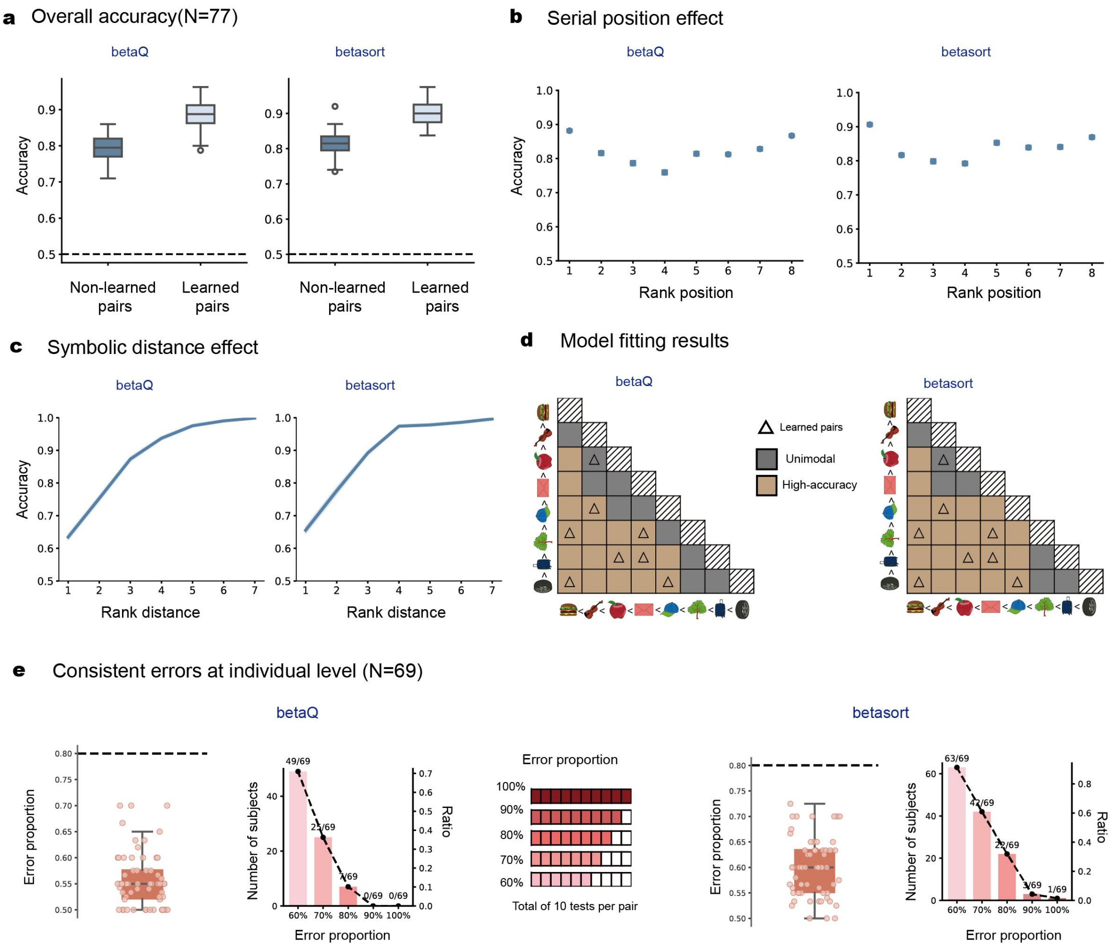


**S4 Fig. BetaQ and Betasort model simulations. (a)** Grand averaged ranking accuracy for learned (dark blue; 8 pairs in few-shot learning) and non-learned pairs (light blue; remaining 20 pairs not directly learned). **(b)** Grand averaged accuracy as function of rank position (serial position effect). **(c)** Grand averaged accuracy as function of ranking distance (distance effect). **(d)** Beta-distribution fitting results for each pair (brown: high-accuracy, $\alpha > 1$, $\beta < 1$; grey; unimodal distribution, $\alpha > 1$, $\beta > 1$). **(e)** Individual-level error consistency based on betaQ (Left) and betasort (Right) models. *Left:* grand averaged error consistency (>0.8, one-sample t-test, $p > 0.05$). dots denote individual data. *Right:* number and proportion of subjects making consistent error on at least one local pair for different thresholds (0.6 to 1.0).

### S5 Figure

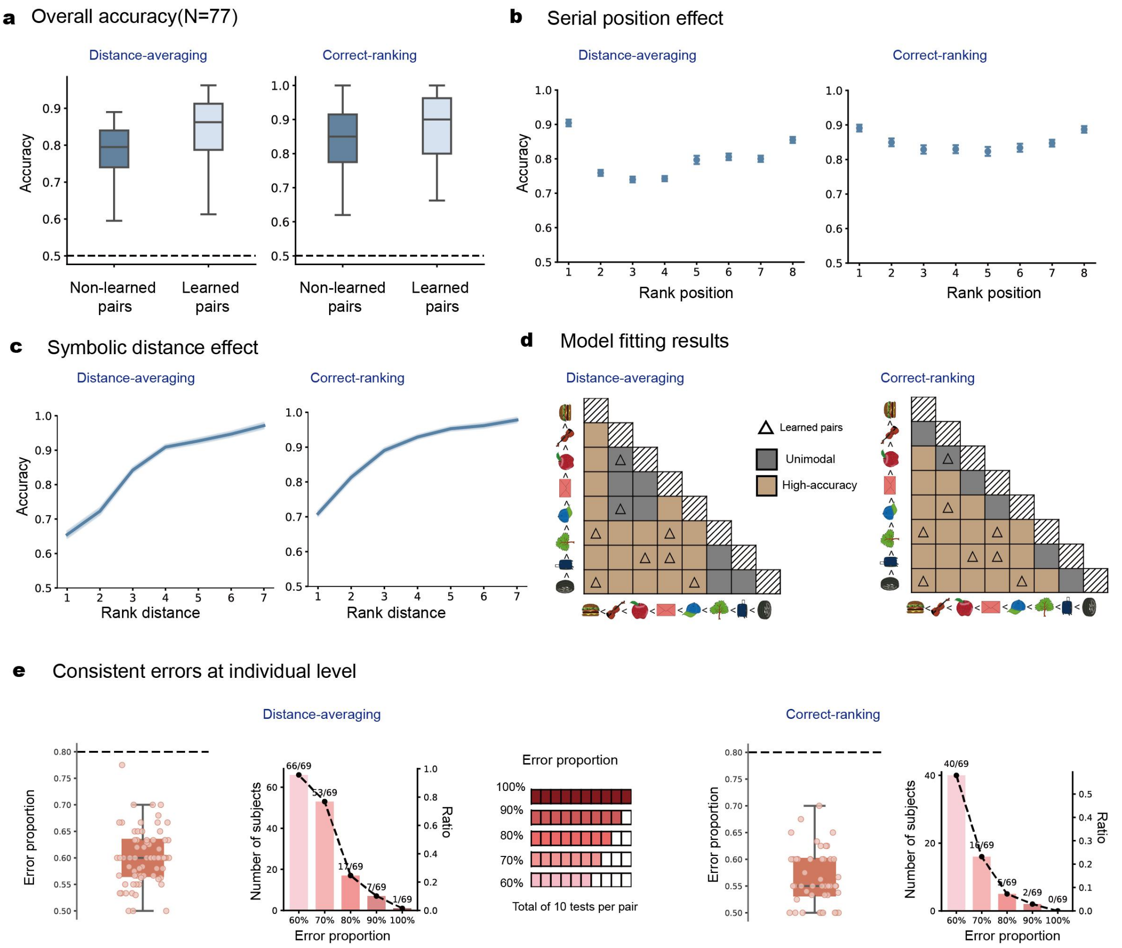


**S5 Fig. Distance-averaging and correct-ranking model simulations. (a)** Grand averaged ranking accuracy for learned (dark blue; 8 pairs in few-shot learning) and non-learned pairs (light blue; remaining 20 pairs not directly learned). **(b)** Grand averaged accuracy as function of rank position (serial position effect). **(c)** Grand averaged accuracy as function of ranking distance (distance effect). **(d)** Beta-distribution fitting results for each pair (brown: high-accuracy, $\alpha > 1$, $\beta < 1$; grey; unimodal distribution, $\alpha > 1$, $\beta > 1$). **(e)** Individual-level error consistency based on Distance-averaging (Left) and correct-ranking (Right) models. *Left:* grand averaged error consistency (>0.8, one-sample t-test, $p > 0.05$). dots denote individual data. *Right:* number and proportion of subjects making consistent error on at least one local pair for different thresholds (0.6 to 1.0).

### S6 Figure

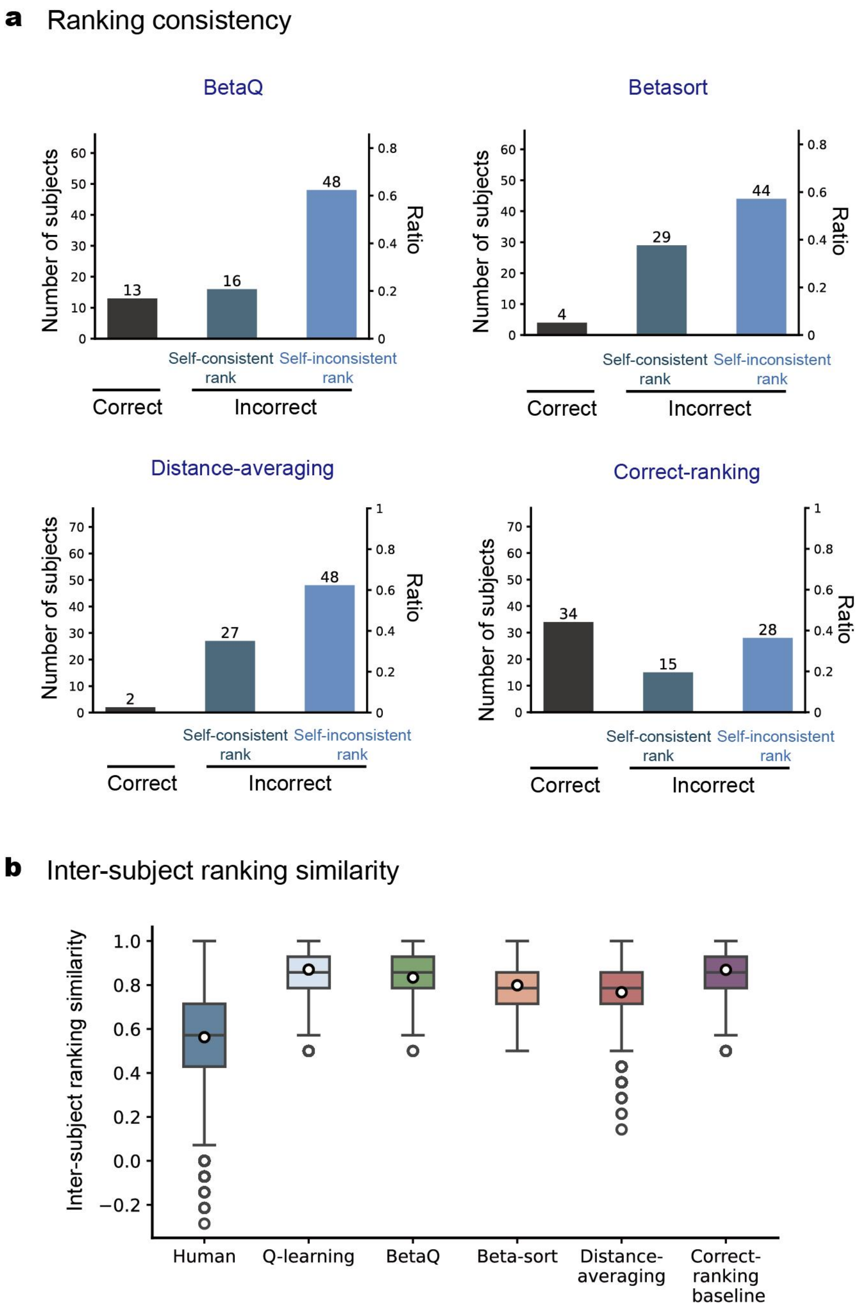


**S6 Fig. Model predicted self-consistency and inter-subject ranking similarity**. **(a)** Number of subjects by self-Consistency category for BetaQ, Betasort, Distance-Averaging, and Correct-Ranking Models. **(b)** Inter-subject ranking similarity for Human, Q-learning, BetaQ, Betasort, Distance-Averaging, and Correct-Ranking Models.

### S7 Figure

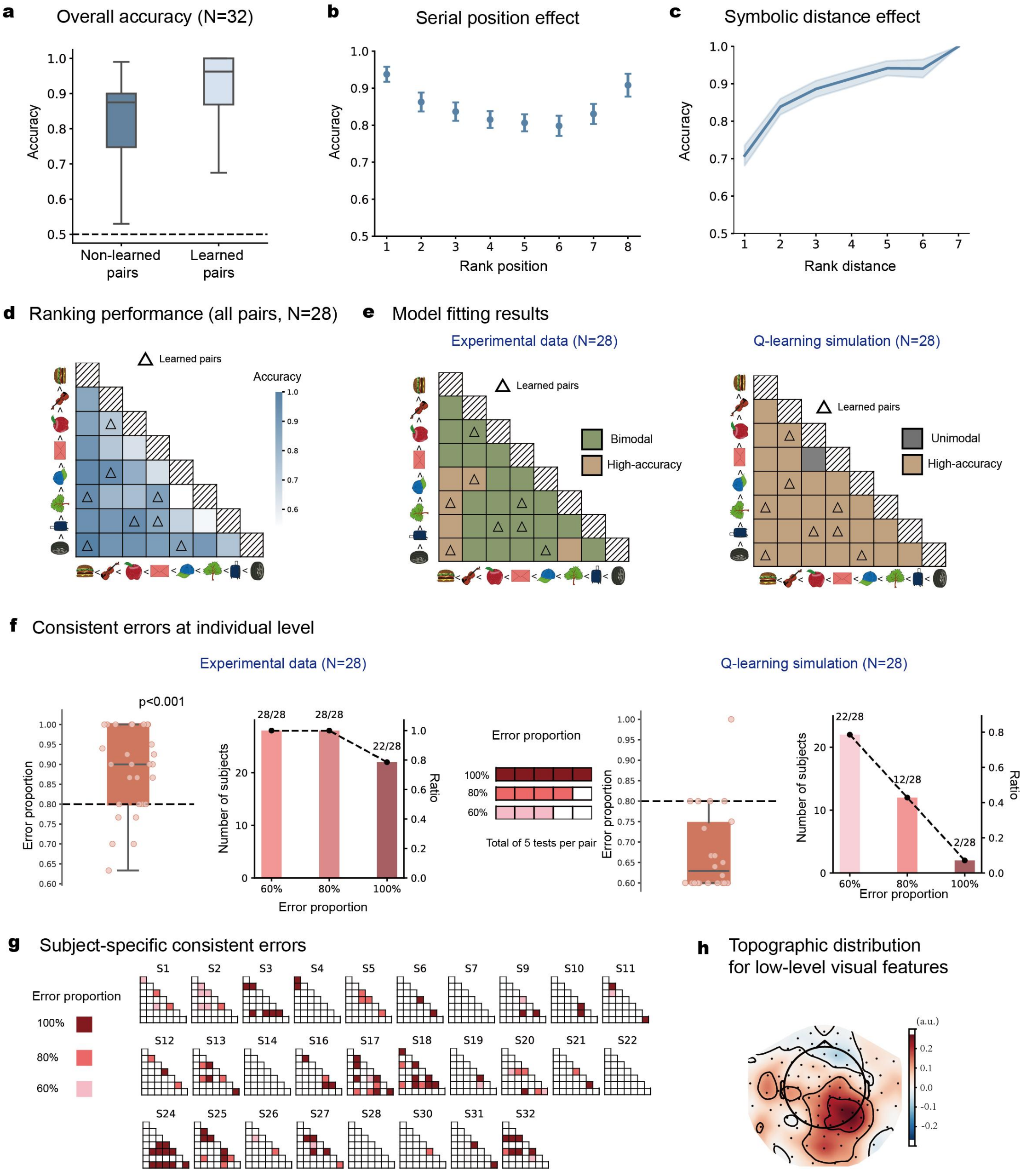


**S7 Fig. Behavioral results and model fitting of MEG experiment. (a)** Grand averaged ranking accuracy for learned (dark blue; 8 pairs in few-shot learning) and non-learned pairs (light blue; remaining 20 pairs not directly learned). **(b)** Grand averaged accuracy as function of rank position (serial position effect). **(c)** Grand averaged accuracy as function of ranking distance (distance effect). **(d)** Grand averaged accuracy matrix for all the 28 tested pairs, with row and column denoting corresponding items per pair (Deep-to-light color represent high-to-low accuracy). Eight directly-learned pairs (few-shot learning) were marked by triangles.
**(e)** Beta-distribution model fitting results for each pair (brown: high-accuracy, $\alpha > 1$, $\beta < 1$; green; bimodal distribution, $\alpha < 1$, $\beta < 1$; grey; unimodal distribution, $\alpha > 1$, $\beta > 1$) for experimental data (left) and Q-learning model (right). Note that simulation

results support unimodal distribution that diverges from bimodal experimental findings. **(f)** Individual-level error consistency and proportion of subjects making consistent error on at least one local pair for different thresholds (0.6 to 1.0), for experimental data (left) and Q-learning model (right). Dots denote individual data. **(g)** Consistent local errors for each subject. **(h)** Topographic distributions for low-level visual features using sensor-level search light analysis.

### S8 Figure

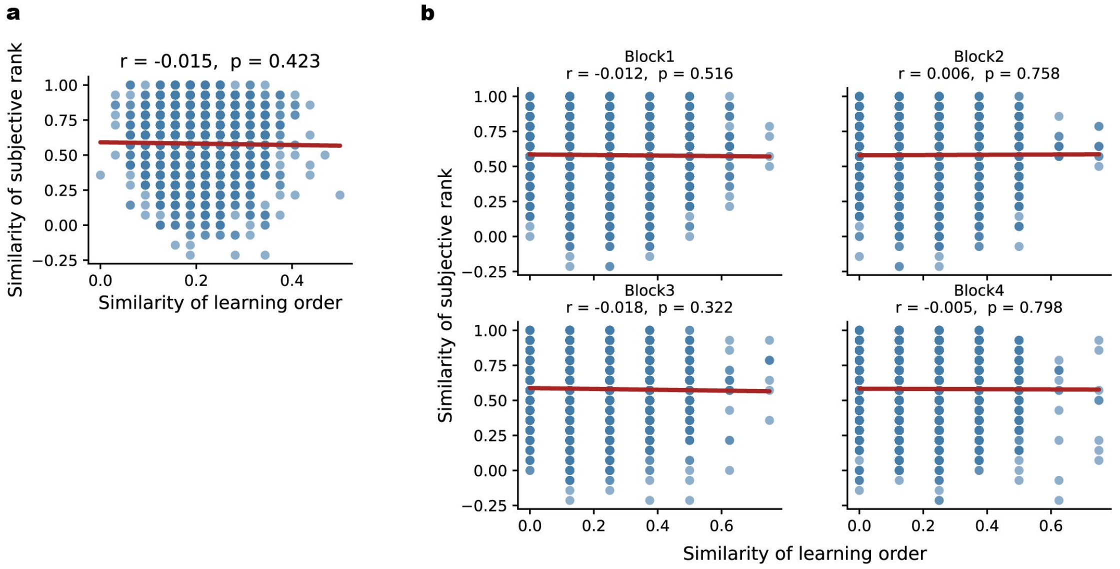


**S8 Fig. Null relationship between learning sequence similarity and ranking similarity. (a)** Overall correlation across all learning blocks revealed no significant association (r = -0.015, p = 0.423). **(b)** Separate analyses for each learning block confirmed the null relationship (Block 1: r = -0.012, p = 0.516; Block 2: r = 0.006, p = 0.758; Block 3: r = -0.018, p = 0.322; Block 4: r = -0.005, p = 0.798).

### S9 Figure

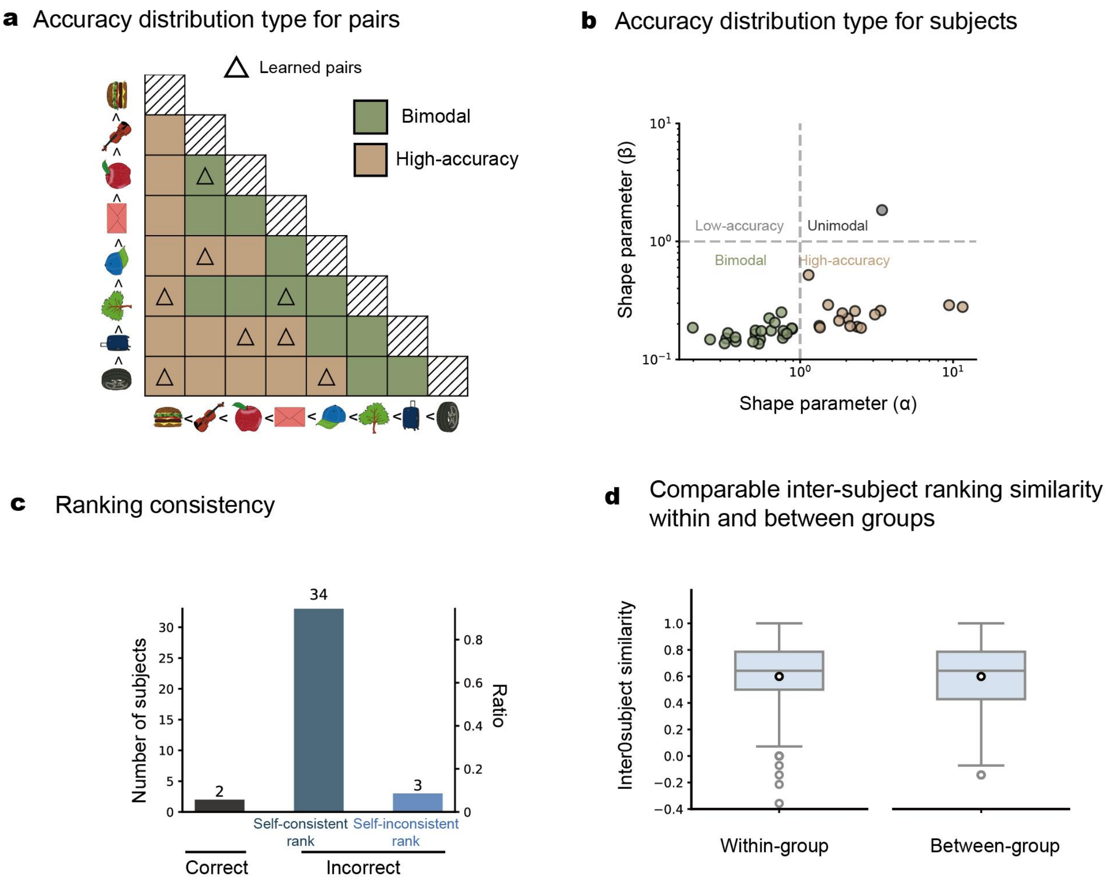


**S9 Fig. Supplemental Between-Group Experiment (N = 40) (a)** Beta-distribution model fitting results for each pair (brown: high-accuracy, $\alpha > 1$, $\beta < 1$; green; bimodal distribution, $\alpha < 1$, $\beta < 1$). **(b)** Beta-distribution model fitting results for each subject. Each point represents the best-fitting α and β parameters for an individual subject. Brown points indicate subjects with bimodal error distributions ($\alpha < 1$, $\beta < 1$); green points indicate subjects exhibiting high-accuracy patterns ($\alpha > 1$, $\beta < 1$). Both α and β are significantly smaller than 1 (one-sample *t* test, excluding 15 high accuracy participant as pre-registered; α, $t(23) = -10.11$, $p < 0.001$; β, $t(23) = -142.26$, $p < 0.001$). **(c)** Number of subjects for each self-consistency category. **(d)** Independent samples t-test revealed comparable ranking consistency among participants who learned the same stimulus set (within-group) versus different stimulus sets (between-group), t= 0.02, p = 0.99. This null result suggests that the specific stimulus set used during learning did not systematically influence participants' subsequent ranking strategies.

**S10 Figure**

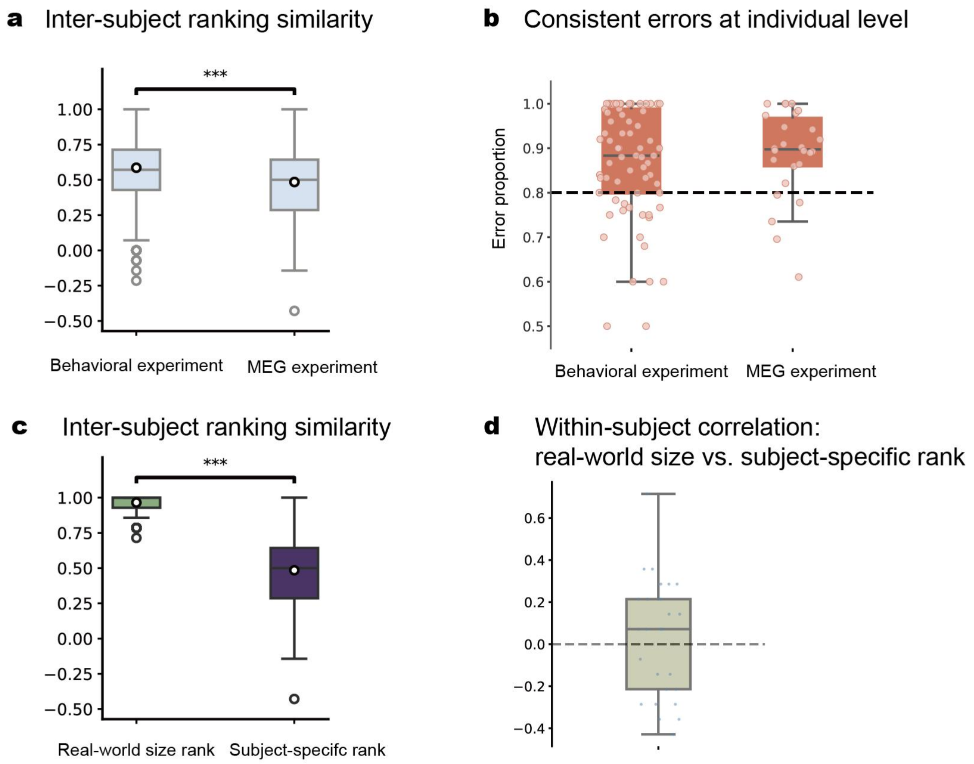


**S10 Fig. Comparison of inter-individual variability and within-subject consistency between behavioral and MEG post-learning phase. (a)** Lower inter-subject ranking similarity in MEG post-learning phase compared to behavioral experiment. **(b)** Consistent errors at individual level in MEG post-learning phase compared to behavioral experiment. **(c)** Lower inter-subject ranking similarity in MEG post-learning phase compared to MEG pre-learning phase. **(d)** Within-subject correlations between real-world size rankings and individually reconstructed subjective rankings were essentially zero (mean r = 0.03, 95% CI [-0.06, 0.13]).

### S11 Figure

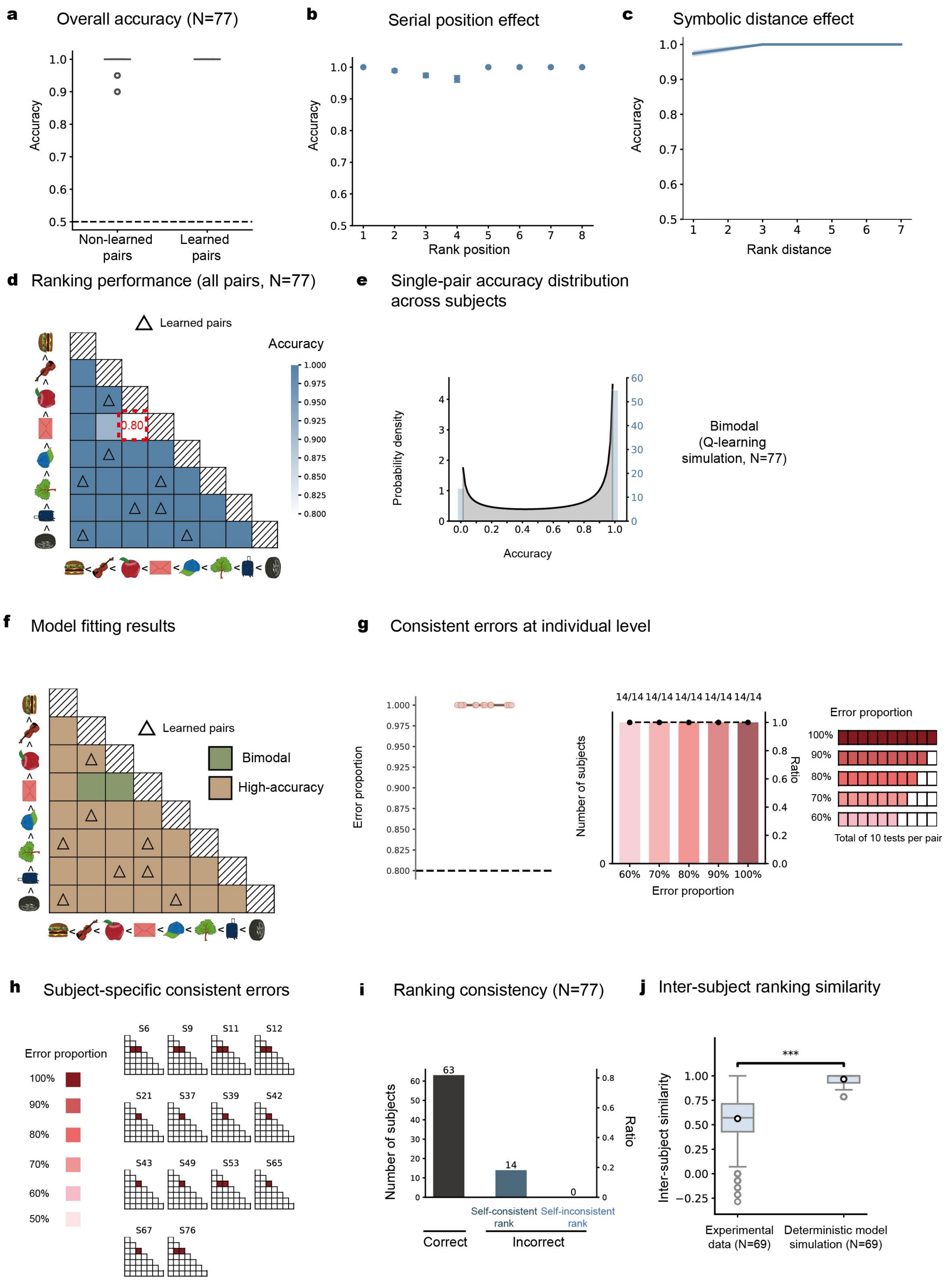


**S11 Fig. Deterministic model simulations. (a)** Grand averaged ranking accuracy for learned (dark blue; 8 pairs in few-shot learning) and non-learned pairs (light blue; remaining 20 pairs not directly learned). **(b)** Grand averaged accuracy as function of rank position (serial position effect). **(c)** Grand averaged accuracy as function of ranking distance (distance effect). **(d)** Grand averaged accuracy matrix for all the 28 tested pairs, with row and column denoting corresponding items per pair (deep-to-light color represent high-to-low accuracy). Eight directly-learned pairs (few-shot learning) were marked by triangles. Red dotted box highlights the exemplar pair in **e. (e)** Observed bimodal distribution, i.e., some subjects consistently made correct inference ($3^{rd} < 4^{th}$), while others made robust errors ($3^{rd} > 4^{th}$).

**(f)**Beta-distribution fitting results for each pair (brown: high-accuracy, $\alpha > 1$, $\beta < 1$; grey; unimodal distribution, $\alpha > 1$, $\beta > 1$). **(g)** Individual-level error consistency based on Distance-averaging (Left) and correct-ranking (Right) models. *Left:* grand averaged error consistency (>0.8, one-sample t-test, $p < 0.05$). dots denote individual data. *Right:* number and proportion of subjects making consistent error on at least one local pair for different thresholds (0.6 to 1.0). **(h)** Local error pattern for each subject. Blank tiles in the lower triangular matrices denote correct pairs (accuracy > 50%). Red tiles denote error pairs (deep-to-light color represent high-to-low error proportion). **(i)** Number of subjects for each self-consistency category. **(j)** Inter-subject ranking similarity for experimental data and Deterministic simulation (two-sample *t*-test, $p < 0.001$).
